# Genomic analyses provide insights into peach local adaptation and responses to climate change

**DOI:** 10.1101/2020.01.15.907709

**Authors:** Yong Li, Ke Cao, Nan Li, Gengrui Zhu, Weichao Fang, Changwen Chen, Xinwei Wang, Xiuli Zeng, Jian Guo, Shanshan Zhang, Qi Wang, Tiyu Ding, Jiao Wang, Liping Guan, Junxiu Wang, Kuozhan Liu, Wenwu Guo, Pere Arús, Sanwen Huang, Zhangjun Fei, Lirong Wang

## Abstract

The environment has constantly shaped plant genomes, but the genetic bases underlying how plants adapt to environmental influences remain largely unknown. We constructed a high-density genomic variation map by re-sequencing genomes of 263 geographically representative peach landraces and wild relatives. A combination of whole-genome selection scans and genome-wide environmental association studies (GWEAS) was performed to reveal the genomic bases of peach local adaptation to diverse climates comprehensively. A total of 2,092 selective sweeps that underlie local adaptation to both mild and extreme climates were identified, including 339 sweeps conferring genomic pattern of adaptation to high altitudes. Using GWEAS, a total of 3,496 genomic loci strongly associated with 51 specific environmental variables were detected. The molecular mechanism underlying adaptive evolution of high drought, strong UV-B, cold hardiness, sugar content, flesh color, and bloom date were revealed. Finally, based on 30 years of observation, a candidate gene associated with bloom date advance, representing peach responses to global warming, was identified. Collectively, our study provides insights into molecular bases of how environments have shaped peach genomes by natural selection and adds valuable genome resources and candidate genes for future studies on evolutionary genetics, adaptation to climate changes, and future breeding.

---

Environmental adaptation is fundamental to species survival and conservation of biodiversity, especially under threats of climate change (Blanquart et al. 2013). Unlike animals, which can escape from hostile environments, plants are sessile and have to adapt by shaping and/or fixing genetic variants that are conducive for survival. Generally, climate is the major selective pressure driving adaptive evolution, resulting in different ecotypes within a single species (Hancock et al. 2011; Fournier-Level et al. 2011). However, the mechanisms underlying how climate shapes plant genomes remain largely unclear. Recently, identifying adaptive variants and understanding molecular mechanism of adaptation across a genome have become tractable due to the advances of sequencing technologies. Recent studies have sought to elucidate genetic bases of adaptation through genome-wide identification of regions under positive selection and/or loci that control adaptive traits in *Arabidopsis thaliana* (Fournier-Level et al. 2011), rice (Yan et al. 2013), sorghum (Lasky et al. 2015), and poplar (Wang et al. 2018). However, no study has focused on genetic bases of adaptation in domesticated perennial fruit crops. Domesticated crops have adapted to diverse climates during domestication and subsequent spread, and show local adaptation through long-term natural selection. Landraces and wild relatives harbor great genetic diversity and an abundance of resistance genes, which provide excellent resources for breeding initiatives. This is especially the case with accessions originating from stressful environments that may have specific stress-resistance genes (Bolger et al. 2014a). However, a cost of domestication is that many resistance related genes have been lost. In addition, global climate change is driving decreases in productivity and distribution changes in several crop species (Tim and Braun, 2013). Therefore, it is of great importance to identify adaptive genes that can contribute to crop improvement, species survival, and global food security in the face of environmental deterioration.

Peach is an important temperate fruit species, with a global yield of 24.7 million tons in 2017 (FAOSTAT; http://www.fao.org/faostat). It is also an important model system for the Rosaceae family, members of which provide one of world’s main resources of fruits. Peach originated in southwestern China, and its landraces and wild relatives are widespread in both temperate and sub-tropical regions, as well as in wet and dry climates (Wang et al. 2012). On the grounds of wide distributions, peach can be regarded as an excellent material for studying adaptation genetics. Peach has a relatively small genome size (∼227.4 Mb) (Verde et al. 2013) and genomic analyses have identified a number of loci and candidate genes associated with human selection and agronomically important traits (Cao et al. 2014; Cao et al. 2016; Li et al. 2019). However, there have been few studies describing genomic loci associated with environmental adaptation and natural selection.

To investigate the genetic basis of local adaptation, we sequenced a wide collection of 263 peach accessions from a broad range of geographical origins and associated with diverse climates, spanning mild and extreme environments. Using the sequencing data, we deciphered adaptive patterns across peach genome by combining the identification of signatures of selective sweeps with genome-wide association studies of environmental variables and adaptive traits. Finally, we identified a candidate gene associated with peach responses to global warming, based on observations over a 30-year period.

## Results and discussion

### Genomic variation map and population structure

We first constructed a genome variation map for peach using a collection of 263 diverse accessions (Fig. 1A), consisting of 52 wild relatives and 211 landraces (Supplementary Table S1), which collectively capture more than 95% of geographic diversity of native distribution of peach landrace and wild relatives. A total of 342.7 Gb of sequence was generated, with a median depth of 5.3 × and coverage of 91.7% of reference peach ‘Lovell’ genome (release v2.0) (Verde et al. 2013) (Supplementary Table S1). We identified a final set of 4,611,842 high-quality single-nucleotide polymorphisms (SNPs) (Supplementary Fig. S1A), of which 1,931,310 were intronic (∼11.33%) and 848,638 (∼4.98%) were exonic. Among SNPs in coding sequence, we found that 7,853 SNPs present in 5,512 peach genes (∼20.5% of total genes) are likely to have a major impact on gene function. The accuracy of identified SNPs was found to be ∼95.6%, based on genotyping of 18 randomly selected SNPs in 130 accessions using a Sequenom MassARRAY platform (Supplementary Table S2). In addition, we identified 1,049,266 small insertions and deletions (INDELs) (shorter than or equal to 6 bp) and 106,388 large structural variations (SVs) (> 30 bp) (Supplementary Fig. S1A).

**Fig. 1.**
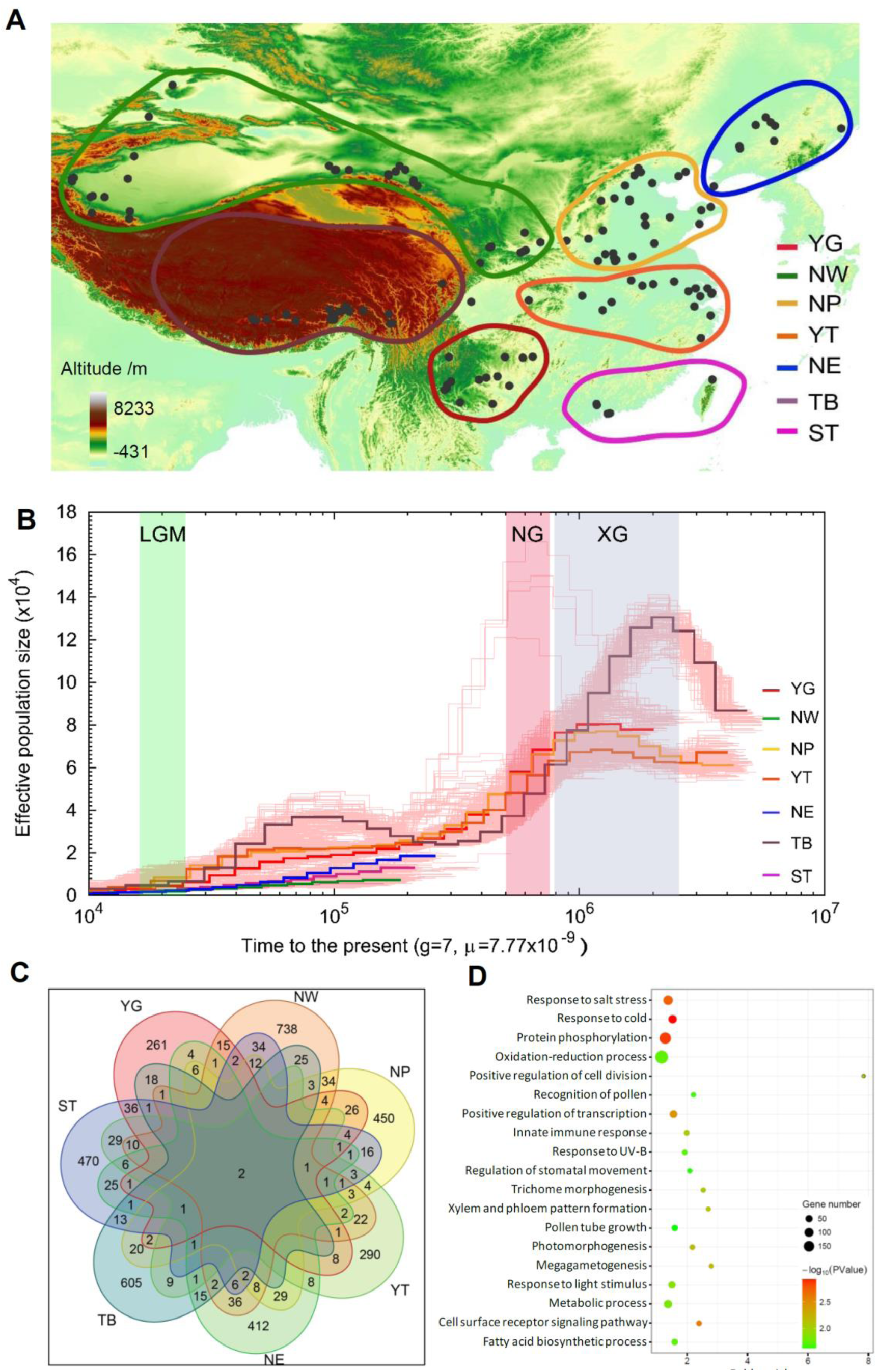
Summary of 263 samples and genes under selection for seven peach groups. (**A**) Geographic distribution of 263 peach accessions used in this study. Each accession is represented by a dot on the world map. Seven ecotypes are highlighted using rings with different colors. (**B**) Demographic history of the seven peach groups. Ancestral population size was inferred using the PSMC model. Three periods of the last glacial maximum (LGM, ∼20 KYA), Naynayxungla Glaciation (NG, 0.5∼0.78 MYA), and Xixiabangma Glaciation (XG, 0.8∼0.17 MYA) are shaded in green, red, and blue, respectively. (**C**) Venn diagram showing the number of genes under selection in the seven groups. (**D**) Over-represented gene ontology (GO) terms in overall selection regions. Only the top 20 most over-represented terms are shown. YG, Yun-gui Plateau. NW, Northwest China. NP, North Plain China. YT, Yangtze River Middle and Backward. NE, Northeast China. TB, Tibet plateau. ST, South China Sub-tropical.

We explored the genetic relationships among 263 accessions using 2,468,307 SNPs with minor allele frequency (MAF) greater than 0.05. Based on the neighbor-joining tree and population structure analysis, the 263 peach accessions could be divided into seven major groups, which were largely congruent with ecotypes classified according to their geographic information, including YG (Yun-gui Plateau), NW (Northwest China), NP (North Plain China), YT (Yangtze River Middle and Backward), NE (Northeast China), TB (Tibet plateau), and ST (South China Sub-tropical) groups (Supplementary Fig. S1B, Supplementary Fig. S2, and Supplementary Table S1). Although the neighbor-joining tree largely supported the division of seven major groups, there were some discrepancies between geographical characterization and phylogenetic clustering (Supplementary Fig. S2D), indicating shared ancestral variation and historical gene flow among landraces in closely related groups. Moreover, principal component analysis (PCA) and model-based clustering analyses also supported the extensive admixture and possible gene flow among landrace groups (Supplementary Fig. S2E and S2F). Furthermore, we found the small pair-wise genetic differentiation (*F*_ST_) values between different landrace groups, again consistent with population admixture (Supplementary Fig. S2G).

Using the demographic analysis with the pairwise sequential Markovian coalescent (PSMC) model (Li and Durbin 2011), we found the sharply decline of effective population size (*Ne*) during the two largest Pleistocene glaciations: the Xixiabangma glaciation (1.17-0.8 MYA) and Naynayxungla glaciation (0.78-0.50 MYA), and a slight decline of *Ne* during the last glacial maximum (∼20,000 years ago) (Fig. 1B).

### Selective sweeps related to adaptation to diverse environments

Peach accessions of each group have adapted locally through long-term selection under local environments (Supplementary Table S3). To identify genomic loci that favor local adaptation for seven groups, we detected signatures of selective sweeps for each group. This revealed a total of 2,092 genomic regions (19.1 Mb, ∼8.4%; 189, 387, 301, 235, 280, 339, and 378 regions for the YG, NW, NP, YT, NE, TB, and ST groups, respectively) (Supplementary Fig. S3), which were termed candidate selection regions (CSRs) (Supplementary Table S4). The overall CSRs harbored 4,198 genes (∼17.5%), including 506, 1,197, 835, 530, 747, 920, and 869 genes for the YG, NW, NP, YT, NE, TB, and ST groups, respectively (Fig. 1C). Selections on these genes may underlie the adaptation to different climates. Notably, we found that few genes were shared among different groups (Fig. 1C), suggesting the unique adaptive patterns for each group and that different climates may shape distinct genomic regions.

We found that genes related to response to different types of stimuli and stress, including temperature, radiation, salt, DNA damage, osmotic, toxin, were overrepresented (*P* < 0.05), suggesting that stress-related genes have participated in adaptive evolution (Fig. 1D, Supplementary Table S5). For instance, two cation/H^+^ exchanger family genes (*CHX*) (*Prupe*.*6G251600* and *Prupe*.*6G251700*) and one *salt overly sensitive 3* (*SOS3*) (*Prupe*.*2G188700*) gene showed high reduction of diversity (ROD) and *F*_ST_ values in the NW group. Homologs of these genes are involved in salt resistance in *A. thaliana* (Monihan et al. 2016), suggesting their potential contributions to adaptation to saline soils in northwestern China. The resistance-related LRR (leucine-rich repeat) domain and PPR (pentatricopeptide repeat) gene family were highly enriched in CSRs (*P* < 0.05) (Supplementary Table S5). The LRR domain, which is considered to be one of the most important domains for plant resistance genes, was also enriched (*P* < 0.05), with 121 of 612 members (19.8%) in CSRs. PPR proteins form one of the largest protein families in land plants that are related to environmental responses, with 286 members in peach genome, of which 79 (∼27.6%) were in CSRs.

The known genes or biological pathways involved in adaptation to the environment in the habitat of each group were determined. For instance, the YG group was distributed on the Yun-gui plateau (Southwest China), a low-latitude and high-altitude (∼2000 m) region with high annual precipitation (> 1100 mm) and acidic soil (pH 4.5∼5.5) (Supplementary Table S3). Genes related to metal ion (including potassium, iron, and zinc) binding and transport, cell membrane function, and response to toxins were overrepresented in this group (107 genes, *P* < 0.05) (Supplementary Table S5), consistent with functions in overcoming cation deficiency and aluminum toxicity that are common in acidic soils (Seguel et al. 2013). For the YT group, we observed enrichments of the LRR domain (24 genes), NB-ARC domain (8 genes), and other genes related to stress responses (32 genes) (*P* < 0.05) (Supplementary Table S5), in comparison to other groups. This suggests that the YT group has accumulated more abiotic and biotic stress-resistance variants due to strong selective pressures in high temperature and high humidity areas (Supplementary Table S3). These results indicate that accessions from the YT group may exhibit higher adaptability than other landrace accessions.

### Genome-wide environmental association studies of 51 environmental variables

Although we obtained candidate genes underlying adaptation by identifying selective sweeps, many adaptive events in natural populations may occur by polygenic adaptation, which would be largely undetected by conventional methods for detecting selection (Pritchard and Di Rienzo 2010). However, local adaptation can generate correlations between environmental variables (EVs) and genomic loci which can be used to detect polygenic adaptation. We investigated a total of 51 EVs of the geographic origin of each accession that are important for plant adaptation (Supplementary Table S6 and S7), representing extremes and seasonality of temperature and precipitation, latitude, altitude, relative air humidity, water vapor pressure, growing season lengths, and radiations. Using a Mantel test, we found a significant correlation between geographic and genetic distances (Pearson’s *r* = 0.73, *P* = 0.000999), with most associations being driven by altitude. To obtain loci associated with EVs, we performed GWEAS on 51 EVs. A total of 9393 association SNPs (Supplementary Table S8), involving 3807 genes, were identified (Fig. 2A). Notably, we found an EV association hotspot regions at the top of chromosome 2 that was enriched with genes encoding NB-LRR proteins in peach genome (Verde et al. 2013). Consistent with the high correlations among some climate variables (Supplementary Fig. S4), only 3496 association SNPs were unique, and ∼62.8% of the associations were shared across different types of EVs, suggesting that different EVs may shape same genomic regions. Notably, a total of 82 genomic loci associated with more than 10 EVs were identified.

**Fig. 2.**
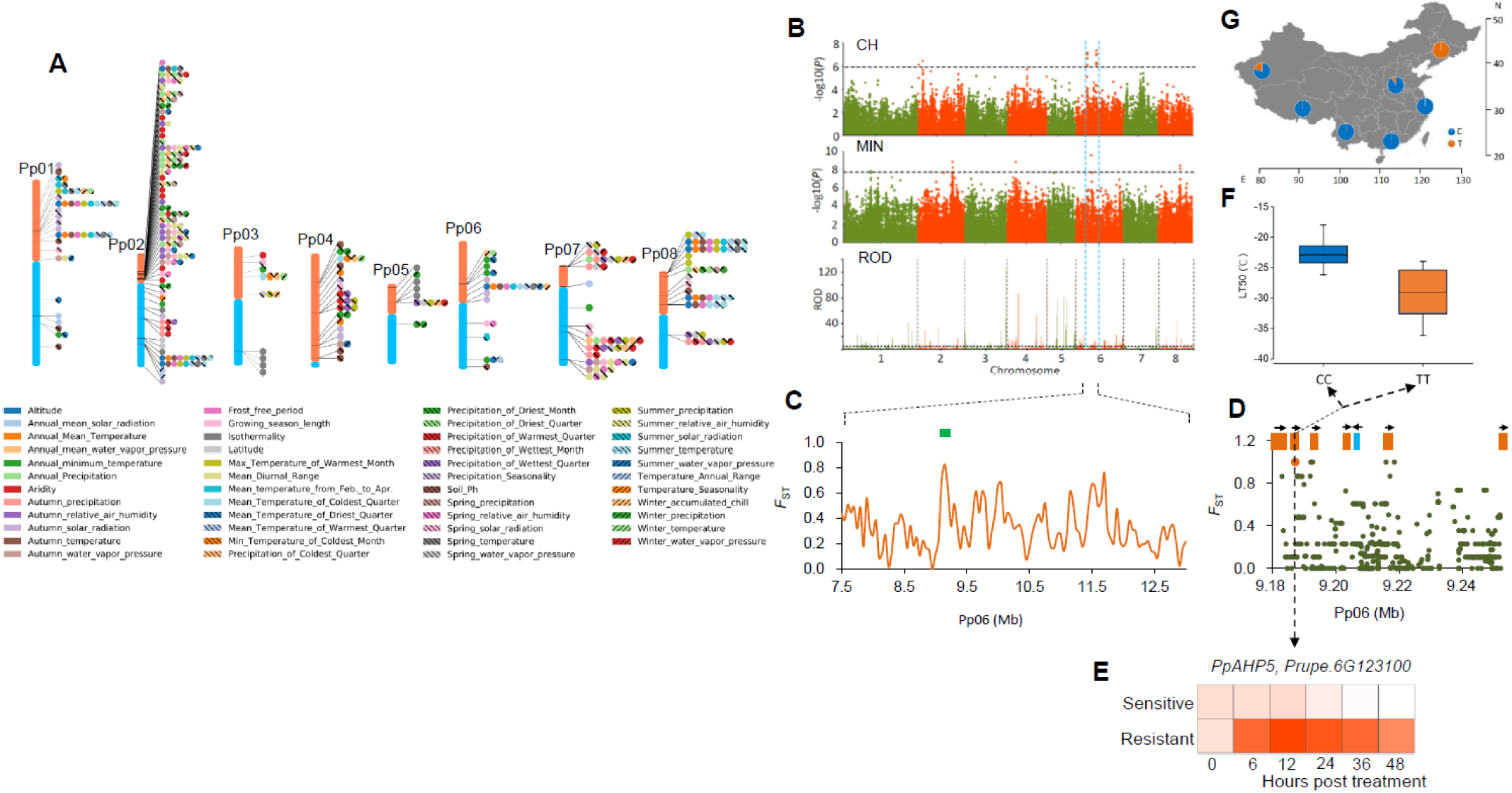
Genome-wide environmental association studies of 51 environmental variables and genomic loci associated with winter cold adaptation. (**A**) SNPs associated with environmental variables (EVs). Only the top 10 association signals for each EV are shown. All signals were included if the total number of signals was < 10. (**B**) The *PpAHP* locus involved in adaptation to winter low temperature in peach. Manhattan plots for a GWAS study of cold hardiness (CH) and winter lowest temperature (MIN), and selection signals of the NE group (ROD) were detailed. The dashed line represents the significance threshold for each test. The candidate genomic region is highlighted between two dashed blue vertical lines. (**C**) Distribution of *F*_ST_ values between NE and ST groups in the candidate region. The green bar indicates the *PpAHP* locus. (**D**) Close-up view of the *F*_ST_ values in a region corresponding to the green bar in (**C**). This region contains six *PpAHP* homologs (orange) and one other gene (light blue). The candidate SNP is highlighted using an orange dot. (**E**) Relative expression changes of *PpAHP5* after cold treatment (−28°C) in resistant and sensitive cultivars. (**F**) Association between genotypes and cold hardiness (lethal temperature of 50%, LT50). (**G**) Allele frequencies of association locus (Pp06: 9,187,362) in *PpAHP5* across seven groups.

Next, we identified known biological processes that were overrepresented among associations for each EV and for overall EVs (Supplementary Table S9). Functional categories related to response to a series of abiotic or biotic stimuli, “programmed cell death (PCD)”, “innate immune response”, and “LRR domain” were highly overrepresented (*P* < 0.05), suggesting that EVs mainly shaped genomic regions related to stress responses. Notably, a series of processes involved in secondary metabolism, including “flavonoid metabolic process”, “jasmonic acid (JA) biosynthetic process”, and “biosynthesis of plant hormones and terpenoids”, were significantly overrepresented (*P* < 0.05) (Supplementary Table S9). We found that genes related to JA biosynthesis were enriched in altitude associations (*P* < 0.05). Previous studies have shown that JA treatment contributes to enhanced cold resistance by promoting expression of the ICE-CBF/DREB1 transcriptional pathway, while a mutation in a key JA biosynthesis gene, *LOX1* (*Prupe*.*6G324400*, an altitude association gene in this study), leads to cold hypersensitive phenotypes (Hu et al. 2013). For each EV, several known biological processes were overrepresented (Supplementary Table S9). For instance, genes involved in ion transport were highly enriched in those associated with soil pH (*P* < 0.05), as soil pH affects absorption of metal ions in plants (Harter 1983).

Temperature and precipitation are two of the most important EVs that affect plant distribution and survival. We identified temperature associated SNPs, distributed across all eight peach chromosomes, and five association hotspots on chromosome 1, 2, 5, 6, and 8 were detected in GWEAS for more than nine temperature-related EVs and altitude (Supplementary Fig. S5A and 5B). Tolerance to low temperature in winter is a major factor that restricts the spread of peach to extremely cold regions (north of 40 °N). To characterize genetic loci underlying adaptation to extremely cold climates in peach, we performed a GWAS analysis of cold hardiness and identified four association peaks, on chromosomes 2, 4, 6, and 7 (Fig. 2B). Of these, the peak on chromosome 6 showed a strong selection signal, with sharp ROD in the NE group that experienced an extreme cold winter (lowest temperature < −30 ^o^C) (Fig. 2B). Moreover, this peak overlapped with the temperature association hotspot on chromosome 6 and association peaks of annual lowest temperature (Fig. 2B). The NE group (*n* = 19) inhabits areas north of 40 °N that have extremely low winter temperatures, while the ST group (*n* = 14) grows in a contrasting climate, south of 25 °N in areas with a warm winter (lowest temperature > 10°C). We searched for genomic regions and SNPs with extremely high differentiation between ST and NE groups. One (Pp06: 9,187,362) of these SNPs (*F*_ST_ = 1) resided within the overlapping intervals between annual lowest temperature and cold hardiness associations (Fig. 2C). This SNP was located in the gene *PpAHP5* (*Prupe*.*6G123100*), belonging to a gene cluster encoding six histidine phosphotransfer proteins (AHP) (Fig. 2D), which have been reported to be involved in mediating cold signaling in *A. thaliana* (Jeon and Kim 2013). Using cold treatment, we found this gene was up-regulated by cold and resistance cultivars harbored significantly high expression level than sensitive one (Fig. 2E). At this SNP locus, all representative accessions in the NE group showed a distinct genotype (TT) compared with the ST group (CC) (Fig. 2F), indicating that the TT genotype in *PpAHP5* is favored in high-altitude cold regions (Fig. 2G), and that *PpAHP5* is a candidate for conferring cold resistance in peach. We also detected six strong association regions for precipitation-related EVs, including annual and seasonal precipitation, length of growing season, aridity, and relative air humidity (Supplementary Fig. S5C and 5D). An extremely strong association hotspot on Pp02 (4.8∼7.2 Mb) was identified, exhibiting enrichments of *R* genes (Verde et al. 2013), *RLKs* super family genes, NB-ARC domains, and other stress response-related genes, suggesting a genetic basis for precipitation adaptation.

To further elucidate the pattern of adaptation, we detected overlaps between selective sweeps and GWEAS. A total of 888 genes (∼23.3% of GWEAS genes) were shared between selective sweeps and GWEAS (Supplementary Fig. S6). This revealed that although selective sweeps are important, adaptations from standing variation or polygenic adaptation are also likely an important mode of adaptation in peach, which may be related to its shortly spread history after domestication (Li et al. 2019). These findings suggest that domesticated fruit species, such as peach, are generating and enhancing adaptation by standing selection on existing multiple sites. This situation is different from *A. thaliana*, which may have reached its adaptive limits owing to the constraints imposed by the limited generation of new mutations (Hancock et al. 2011). Collectively, these results indicate that both selective sweeps and GWEAS are central factors in the adaptive genetics of domesticated species.

### Adaptation to highly drought regions

The NW group is from northwestern China, which has an extreme climate, characterized by severe aridity (< 150 mm annual rainfall) (Fig. 3A) and extreme high or low temperatures in the summer (> 40 °C) or winter (< −30 °C) (Supplementary Table S3). Peach accessions from this region are frequently challenged by high drought stress. We found that genes overrepresented in this group included those involved in abscisic acid (ABA) biosynthesis and signal transduction (*P* < 0.05) (Supplementary Table S5), which are well known to regulate drought stress responses. Transcriptome analyses of peach accessions grown under drought stress conditions revealed that genes involved in the ABA pathway were highly enriched among differentially expressed genes (DEGs), including *NCED, PYR, ABA2, PP2C*, and *ABRE* genes that showed selective signals in the NW group (Fig. 3B), further suggesting a key role of ABA pathway in peach drought responses.

**Fig. 3.**
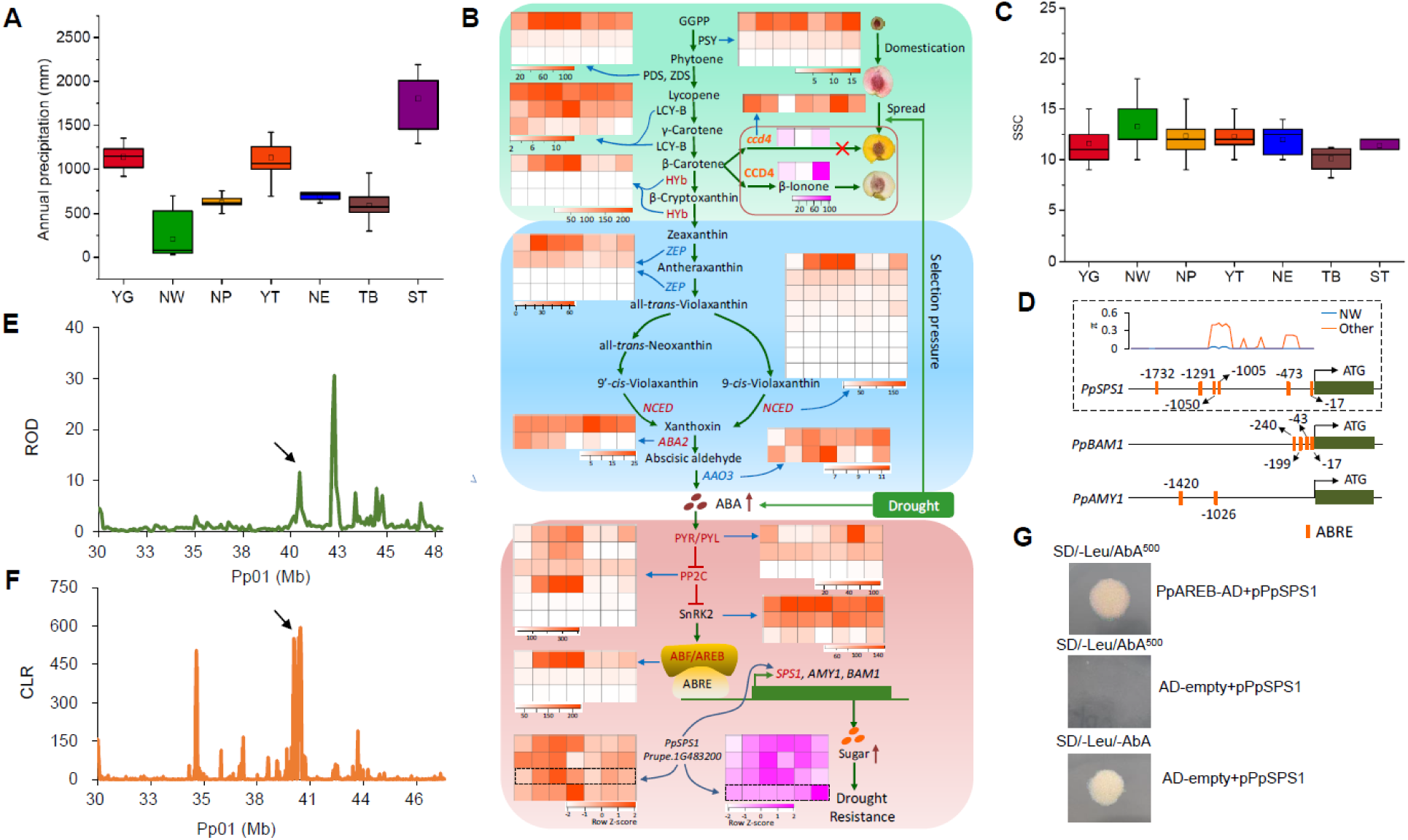
Genetic basis of drought resistance and high sugar content in the NW group. (**A**) Annual precipitation among the seven groups. (**B**) Relationship between the ABA pathway, drought stress and evolution of flesh color. Heat map in orange indicate gene expression levels (FPKM) under drought stress (0h, 6h, 12h, 24h, 3d, 6d, 12d). Heat maps in pink indicate gene expression levels (FPKM) during peach fruit development (10, 50, and 90 days post bloom date (dpb) for *PpCCD4*; 20, 40, 60, 80, 100, 120 dpb for *PpSPS1*). Genes under selection in the NW group are highlighted in red. Red arrows indicate the increase in levels of ABA and sugars. (**C**) Soluble solid content (SSC) among the seven groups. (**D**) ABRE *cis*-acting elements in the promoters of *PpSPS1, PpBAM1*, and *PpAMY1*. Orange boxes indicate ABRE elements in the promoter of each gene. The number around each ABRE represents the position from the ATG. The distribution of ABRE elements and nucleotide diversity in the promoter of *PpSPS1* in the NW and other groups are shown in a dashed box. (**E**) Distribution of ROD around *PpSPS1* on chromosome 1. Black arrow points to *PpSPS1*. (**F**) Distribution of CLR values around *PpSPS1* on chromosome 1. Black arrow points to *PpSPS1*. (**G**) Verification of the interaction between *PpAREB* (*Prupe*.*1G434500*) and the promoter of *PpSPS1* (*Prupe*.*1G483200*) using a yeast one-hybrid assay.

Sugars function as the important signaling molecules in response to a range of abiotic and biotic stresses in plants (Lastdrager et al. 2014). We found that peach fruits produced by accessions from the NW group, especially accessions from Xinjiang province (Wang et al. 2012), consistently had higher soluble sugar contents than those from other groups (Fig. 3C). Associated long-term natural selection pressures contributing to greater accumulation of soluble sugars likely include aridity, high diurnal temperature variation, and long sunshine duration. Moreover, the starch and sucrose metabolism pathways were overrepresented in both DEGs under drought stress treatment (35 genes) and genes under selection in the NW group (12 genes) (*P* < 0.05), congruent with roles of sugars in drought stress. Furthermore, all the 12 genes in the selective sweeps were differentially expressed following the drought stress treatment. We conclude that higher soluble sugar contents in accessions from northwestern China represent an adaptive trait driven by the local drought environment.

Previous studies of apple have demonstrated that drought stress and ABA contributed to soluble sugar accumulation through the activation of sugar transporter and amylase genes by the ABA-responsive transcription factor, *AREB2* (Ma et al. 2017). Similarly, both drought stress and exogenous ABA induce an increase in soluble sugar accumulation in peach fruit (Kobashi et al. 2000; Kobashi et al. 2001). Here we found that two putative gene targets of AREB2 (Fig. 3B and 3D), *PpAMY1* (*Prupe*.*1G142400*) and *PpBAM1* (*Prupe*.*1G053800*), were up-regulated by drought treatment; however, neither exhibited a significant selection signal. To identify additional target genes in drought mediated sugar accumulation, we searched for genes harboring the putative binding domain of AREB2 among genes under selection in the NW group. This revealed a sucrose phosphate synthase gene (*PpSPS1, Prupe*.*1G483200*), with six ABA-responsive elements (ABREs) in the promoter region (Fig. 3D), showing a strong selection signal, with high ROD and CLR values (Fig. 3E and 3F). *PpSPS1*, which is involved in the biosynthesis of sucrose, the predominant soluble sugar in mature peach fruit and the key factor conferring sweetness, was up-regulated by drought treatment (Fig. 3B), suggesting its roles in drought stress response. The expression of *PpSPS1* increased by ∼500-fold during fruit maturity (Fig. 3B), implying its roles in fruit ripening and sugar accumulation. Using a yeast one-hybrid experiment, we verified the interactions between AREB/ABF and the promoter of *PpSPS1* (Fig. 3G), providing new insight into ABA-mediated enhanced sugar accumulation under drought stress. The selection on sugar related genes may mediate adaptation to drought stress in the NW group, accompanied by the increases in fruit sugar content. In addition, we found that the top of chromosome 5 and the middle of chromosome 4, which have been reported to harbor major SSC-and sugar content-associated quantitative trait loci (QTLs) and SSC candidate gene *PpNCED3* (Martínez-García et al. 2013; Li et al. 2019), also showed strong selection signals in the NW group. Selections on these genes may underlie the genetic basis of high sugar levels in peach accessions grown in areas with high drought stress. Moreover, such genes represent excellent candidates for high-sugar breeding.

Intriguingly, we found that flesh color of peach showed strong geographic pattern, with ∼80% of yellow-fleshed peach landraces originating from northwestern China (NW group). Yellow flesh of peach mainly depends on the content of carotenoids at maturity, including β-cryptoxanthin and β-carotene, and carotenoids are believed to be the major precursors for ABA biosynthesis (Fig. 3B). A previous study has identified three loss-of-function variants involved in a carotenoid cleavage dioxygenase gene (*PpCCD4, Prupe*.*1G255500*), leading to the abnormal carotenoid degradation and yellow flesh (Falchi et al. 2013). The disturbed function of *PpCCD4* in yellow-fleshed peach resulted in the retention of carotenoids, which can provide more precursors for ABA biosynthesis (Fig. 3B), and may contribute to higher ABA levels and subsequent enhanced drought tolerance. Moreover, using transcriptional analyses, we found that *PpCCD4* was down-regulated by drought treatments (Fig. 3B), suggesting its response to drought stress. Furthermore, the carotenoid biosynthetic pathway was highly overrepresented with genes under selection in the NW group (*P* < 0.05). Therefore, we conclude that yellow peach flesh has undergone long-term adaptive selection, driven by drought stress, and that modern yellow-fleshed peach cultivars may originate from northwestern China.

Collectively, we constructed a joint pathway for drought adaptation evolution in peach, driven by the complicated interactions between carotenoids, ABA, and sugar, of which ABA may be the central controller and play the key roles.

### Adaptation to high altitudes

Members of the TB group (*n*=45) are from ‘the roof of the world’, Tibet plateau, which is the highest plateau on the earth, with an average elevation of 4500 m. This area is inhospitable to many organisms because of its strong ultraviolet radiation, hypoxia, and severe cold (Supplementary Table S3). At high altitudes, genome integrity is continuously challenged by intensive solar ultraviolet radiation (UV-B, 280-315 nm)-induced DNA damage. Peach accessions in the TB group tolerate these conditions using several adaptation-related phenotypes, such as a dark branch color, epigeal germination, and red-colored new shoots (Supplementary Fig. S7). We identified 339 genomic regions, harboring 920 genes, showing signals of natural selection in the TB group (Supplementary Table S4). Of which, we found a significant enrichment of genes involved in ‘response to UV-B’ category (*P* = 0.0004) (Supplementary Table S5), which is consistent with the adaptation to high-altitude origin of the TB group. Flavonoids are a group of plant secondary metabolites, which play important roles in UV-B protection (Li et al. 1993), and we found two genes in the flavonoid biosynthetic pathway in the ‘response to UV-B’ category (Fig. 4A): chalcone synthase (*PpCHS2, Prupe*.*4G252100*) and phenylalanine ammonia-lyase (*PpPAL, Prupe*.*6G235400*), both of which showed strong selection signals in the TB group, with high μ and Tajima’s *D* values (Fig. 4B and 4C). Chalcone synthase catalyzes the first committed step in flavonoid biosynthesis and previous studies showed that functional perturbation of an *A. thaliana* homolog, *AtCHS*, resulted in UV-hypersensitive phenotypes, while in a UV-B-tolerant mutant *Atchs* was up-regulated (Birza et al. 2001). We found that *PpCHS2* was highly and specifically expressed in the phloem of new shoots in the TB group (Fig. 4D), consistent with the red new shoot phenotype. By scanning genomic variants in or around *PpCHS2*, we found that a SNP (Pp04: 16,896,126, A>T) causing the introduction of a premature termination codon (Fig. 4E) showed a high frequency in low altitude accessions (76.3%), but extreme low frequency of substitution allele in the TB group (2.0%). Intriguingly, this SNP was located at the key active region for protein function, CoA-binding motif (Fig. 4F), leading to an incomplete binding motif that may result in the loss of function. Moreover, the premature termination resulted in the loss of one conserved catalytic residue which is also crucial for catalytic activity (Ferrer et al. 1999). Therefore, this SNP was designated as a candidate causative variant for the red new shoot phenotype involved in flavonoid-mediated UV-B adaptation. Collectively, our results suggest that selection on *CHS* gene and the regulation of anthocyanins may be one of important mechanisms to confer avoiding damage from UV irradiation for peach at high altitudes.

**Fig. 4.**
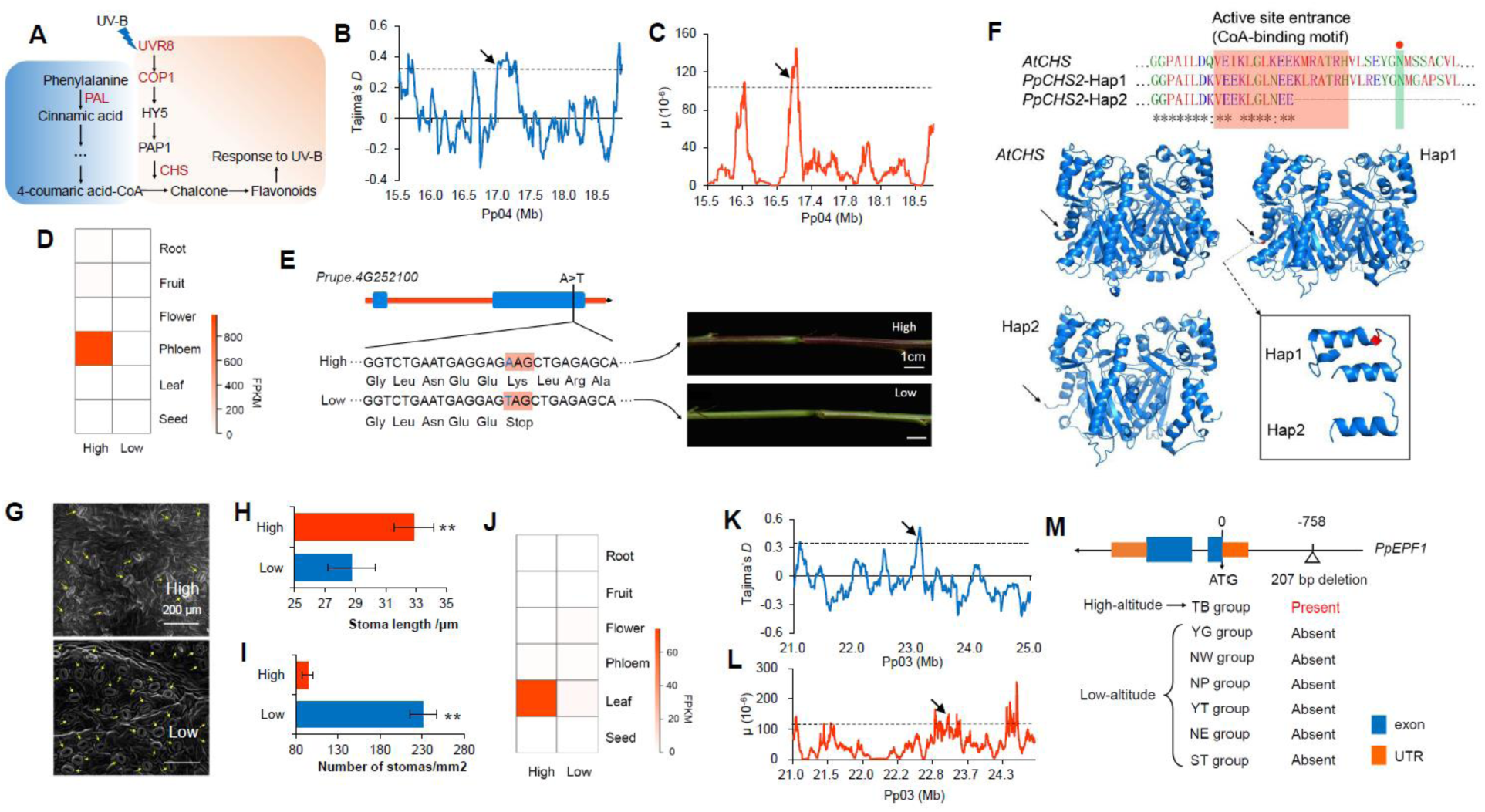
Genomic regions and candidate genes related to high-altitude adaptation in Tibet. (**A**) Pathway related to plant response to UV-B. Genes under selection are highlighted in red. (**B-C**) Distribution of Tajima’s *D* (**B**) and μ values (**C**) in the region around *PpCHS2* (*Prupe*.*4G252100*) on chromosome 4 (15.5-19.0 Mb). The dashed horizontal lines indicate a threshold of top 5% for Tajima’s *D* (≥ 0.36) and μ test (≥ 1.07). Arrows point to *PpCHS2*. (**D**) Heatmap of expression profiles of *PpCHS2* in different tissues in low- and high-altitude accessions. (**E**) A candidate stop-gained SNP in *PpCHS2* that is associated with high altitude adaption and new shoot colors in accessions from low- and high-altitudes. (**F**) Effects of stop-gained SNP on protein structure of CHS. 3D structure of CHS protein was obtained from Swiss-prot. The red shadow represents the CoA-binding motif. The green shadow represents one of the conserved enzyme active site. (**G**) Scanning electron microscopy (SEM) of stomata from the leaves of high- and low-altitude accessions. The magnification is 800×. (**H-I**) Stomatal length (**H**) and stomatal density (**i**) in high- and low-altitude accessions. ** indicates *P* < 0.01. (**J**) Heatmap of expression profiles of *PpEPF1* in different tissues in accessions from low- and high-altitudes. (**K-L**) Distribution of Tajima’s *D* (**K**) and μ values (**L**) in a region around *PpEPF1* (*Prupe*.*3G235800*) on chromosome 3 (21.0-25.0 Mb). The dashed horizontal lines indicate a threshold of top 5% for Tajima’s *D* (≥ 0.36) and μ test (≥ 1.07). Arrows point to *PpEPF1*. (**M**) Structure of *PpEPF1* and the position of the 207-bp deletion. The presence and absence of the 207-bp deletion in the seven groups are given.

We observed that, compared with low-altitude accessions, those from high-altitudes had a lower density of stomata and larger stomata size (Fig. 4G-4I). This may represent an adaptive evolution to hypoxia at high altitudes. Interestingly, we found that the biological category ‘stomatal complex patterning’ was significantly enriched in the gene set under selection (*P* = 0.008). By transcriptional analyses of these genes, we found one of them, *Prupe*.*3G235800*, was highly and specifically expressed in leaves, showing an altitudinal pattern with higher expression levels in the TB group than in the low-altitude group (Fig. 4J). Notably, *Prupe*.*3G235800*, which encodes the epidermal patterning factor 1 (PpEPF1) involved in stomatal development (Hara et al. 2009), showed strong selection signals, based on the high Tajima’s *D* and μ values (Fig. 4K and 4L). Previous studies have shown that the mutation of a homolog of *PpEPF1* in *A. thaliana* results in increased stomatal density (Hara et al. 2009). By scanning the variants in *PpEPF1*, we found that SNPs with functional significance were absent. Through further scanning variants at the upstream or downstream of *PpEPF1*, we identified a TB group specific 207-bp deletion in the promoter region (−758 bp from the start codon) of *PpEPF1* (Fig. 4M), suggesting that the adaptive evolution controlled by *PpEPF1* may be mediated by regulation of its expression. Furthermore, over-expression of *PpEPF1* in *A. thaliana* resulted in a decrease in stomatal density (Supplementary Fig. S8). These findings suggest that selection on *PpEPF1* may be closely related to adaptation to hypoxia in high-altitudes through the regulation of stomatal density.

### A major *SVP* locus involved in adaptive evolution of bloom date

Bloom date (BD) is crucial for local adaptation in peach, and is controlled by multiple genes (Fan et al. 2010). To explore the genetic basis of adaptation of BD, we performed GWAS of BD using 174 accessions that were phenotyped. This revealed 399 associated SNPs and 12 association peaks (Fig. 5A), of which six overlapped with previously reported QTLs (Fan et al. 2010). Next, we identified candidates involved in local adaptation by detecting SNPs showing associations with EVs using a latent factor mixed-effect model (LFMM), resulting in a final set of 23 association peaks (Fig. 5A). By overlapping BD GWAS and LFMM analyses, we found four regions on chromosomes 3, 5, 6, and 8 that may underlie the local adaptation of BD during spread of peach to different climates (Fig. 5A).

**Fig. 5.**
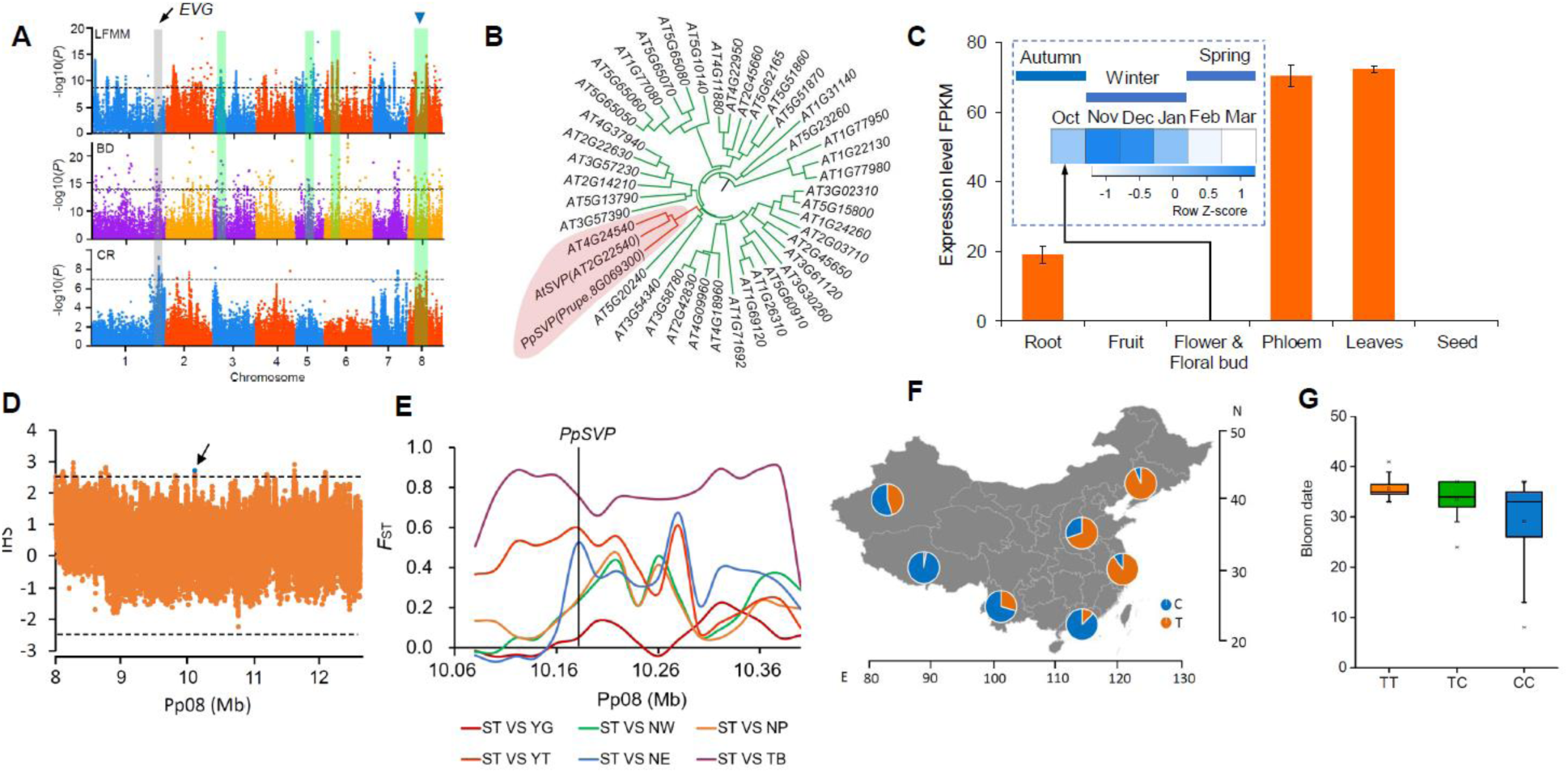
A major *PpSVP* locus involved in local adaptation of bloom date in peach. (**A**) Manhattan plots of SNPs associated with EVs (LFMM), bloom date (BD), and chilling requirement (CR). Dashed lines represent the significance thresholds for the tests. The overlapped regions between GWAS for BD and LFMM are highlighted using green shaded rectangles. The major QTL for CR and BD overlapping with local selection signals on chromosome 8 surrounding *PpSVP* is indicated by a blue triangle. The *EVG* locus is highlighted using a gray shaded rectangle. (**B**) Neighbor-joining tree of *PpSVP* and MIKC-type MADS family genes. The clade containing *PpSVP* is highlighted in red. (**C**) Temporal and spatial expression patterns of *PpSVP*. Error bars represent standard deviation of three biological replicates. (**D**) Patterns of normalized iHS scores across the ∼4 Mb genomic region around *PpSVP*. The dashed horizontal lines represent the threshold of positive selection signal (|iHS| > 2.5). The blue dot indicates the SNP (Pp08: 10,173,576) that showed high iHS score in *PpSVP*. (**E**) *F*_ST_ around *PpSVP* among different groups. The associated SNP in *PpSVP* is indicated using vertical black line. (**F**) Allelic frequencies of the associated SNP (Pp08: 10,173,576) in *PpSVP* across seven groups. (**G**) Relationship between genotypes of associated SNP (Pp08: 10,173,576) and bloom date.

Chilling requirement (CR) is another important adaptive trait and is significantly correlated with BD. We re-performed the GWAS for CR based on our previous study (Li et al. 2019) using 174 landrace accessions and identified six association peaks, of which three (chromosome 1, 7, and 8) were shared with BD (Fig. 5A), including the major QTL for CR harboring the *EVG* locus conferring dormancy mutation in peach (Li et al. 2009). After overlapping GWAS of CR and BD with the LFMM analysis, we found a strong overlap spanning ∼4-Mb on chromosome 8, which may be important for local adaptation of BD in peach (Fig. 5A). Interestingly, the major QTL for CR and BD on chromosome 1 showed no local adaptation signal in the LFMM analysis (Fig. 5A), suggesting that climates may drive the evolution of BD and CR by shaping QTLs with small effects.

The 4-Mb region encompasses 275 genes, including a putative ortholog of *A. thaliana SHORT VEGETATIVE PHASE* (*PpSVP, Prupe*.*8G069300*). *SVP* is involved in controlling flowering time and has previously been implicated in regulating dormancy in *Prunus* (Li et al. 2009; Sasaki et al. 2011; Zhang et al. 2012). Phylogenetic analysis confirmed that *PpSVP* belongs to a MADS-box family and is closely related to the *AGL22* subfamily (Fig. 5B). *PpSVP* showed strong tissue-specific expression, with high expression only in vegetative organs. Moreover, expression of *PpSVP* was up-regulated during dormancy induction and down-regulated by winter chill (0-7.2°C) and by forcing temperature (heat) in floral buds in spring (Fig. 5C), suggesting its potential role in regulating BD and CR. Moreover, through calculating the standardized integrated haplotype score (iHS) for SNPs located in this overlap region, we found a strong positive selection signal around the *PpSVP* locus (Fig. 5D). Additionally, an exceptionally high *F*_ST_ value was identified in this region, especially between the ST and NE groups and between the ST and YT groups (Fig. 5E) that harbor distinct bloom date. The *PpSVP* locus thus represents a strong candidate gene for local adaptation of BD and CR. We propose that spatially varying selection has driven latitudinal differentiation at this locus. Positive selection signals, revealed by a CLR test, were also detected in the NE and ST groups (Fig. 5F). Overall, all these results provide compelling evidence of local selection on the *PpSVP* locus during adaptive evolution to different climates after domestication.

To identify the causal variants underlying adaptation of BD, we screened for SNPs with high *F*_ST_ between the NE (late bloom) and ST (early bloom) groups at the *PpSVP* locus. No SNP with high differentiation was identified that caused an amino acid change. However, a SNP located at 5’-untranslated regions (5’-UTR) with high *F*_ST_ value (*F*_ST_=0.9) was identified, suggesting that the BD and CR may adapt to different climates through shaping the expression of the controlled gene. Allele frequencies of this SNP showed strong geographical pattern and the early bloom alleles (CC) mainly occurred in low altitude regions (ST and YG groups) and wild group (TB group) (Fig. 5G and 5H), consistent with phenotype. This also provides insights into two distinct evolutionary routes of BD and CR to low and high chill regions. Moreover, overexpression of the low-altitude favored genotype of *PpSVP* (CC) in *A. thaliana* resulted in plants with strong vegetative growth and delayed flowering time (Li et al. 2019).

### Genomic locus associated with response to climate change

Adaptation to accelerating rates of climate change is increasingly important for species survival. The advance in bloom date (ABD), as a consequence of global warming over recent decades, has been observed in many temperate species, including peach (Menzel et al. 2006; Li et al. 2016). However, the genetic mechanism underlying ABD have not been characterized. We performed a long-term observation of BD with 89 peach accessions spanning three decades, from the 1980s to 2010s (Supplementary Fig. S9A). We observed a significant ABD (*P* < 0.001), based on an additive main effects and multiplicative interaction (AMMI) analysis (Annicchiarico, 1997), and the main driver was found to be a temperature rise in the spring (explained 61.3% variation, *P* < 0.001) (Fig. 6A). Using a linear regression analysis, we quantified ABD and found that BD has advanced by approximately 10 days on average over last 30 years (Fig. 6B).

**Fig. 6.**
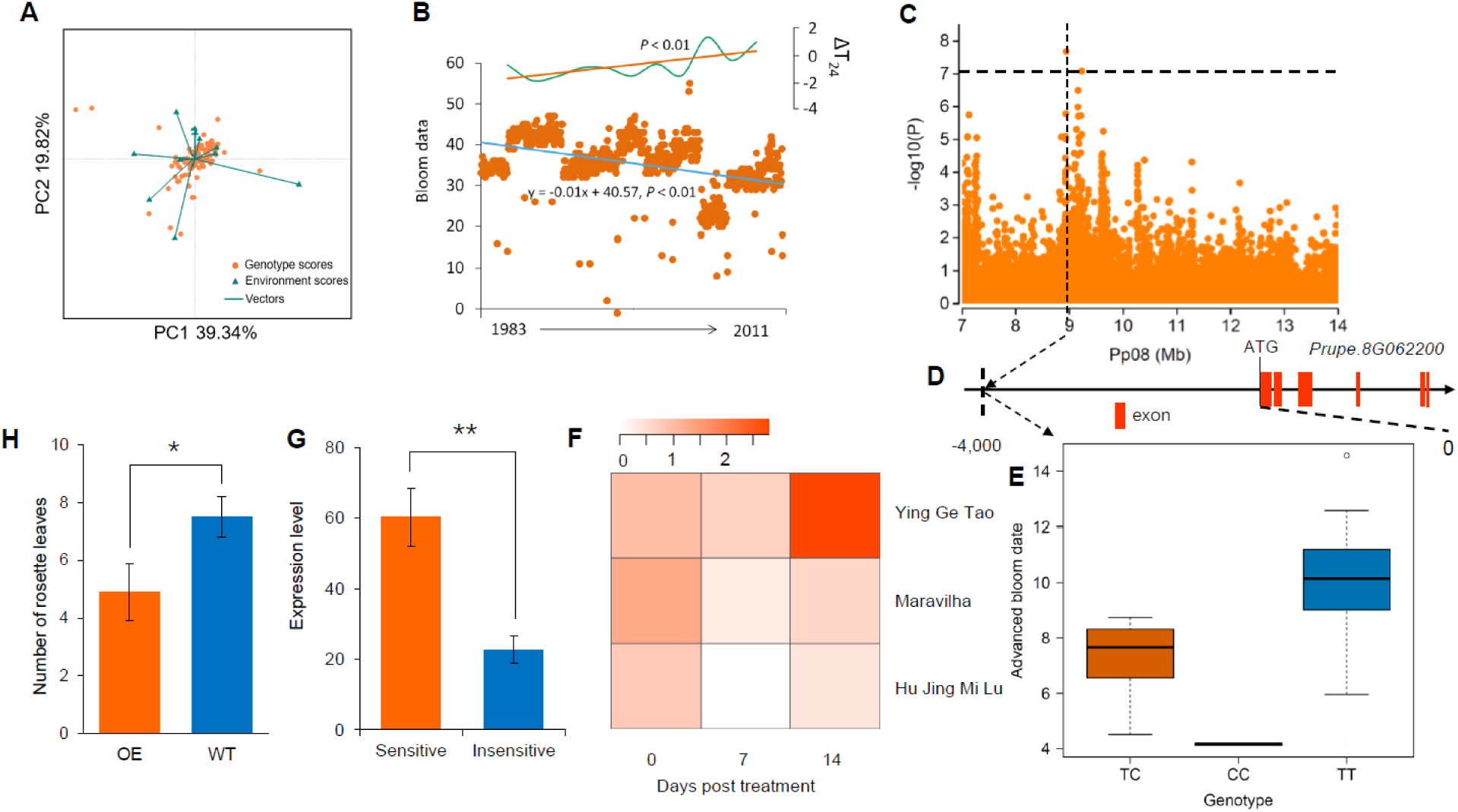
Genotype-environment interaction analysis and genome-wide association study of advance in bloom date. (**A**) Genotype-environment interaction analysis of bloom date from 1983 to 2011 using the AMMI analysis. (**B**) Scatter plots of relative bloom date of 89 peach accessions from 1983 to 2011 and temperature change in the spring. The blue and orange lines represent the trend of bloom date changes and temperature changes in the spring, respectively, based on the linear regression analyses. ΔT24 indicates anomalies in the mean temperature from February to April compared to those from 1983-2011. (**C**) Regional Manhattan plot of GWAS for ABD on chromosome 8 of the 7.0-14.0 Mb region. The gray dashed line indicates significance threshold (*P* < 7.28×10^−8^ or −log10(*P*) > 7.08) using a Bonferroni test (0.05). (**D**) Most significant SNP associated with ABD and its location relative to gene *PpLNK1* (*Prupe*.*8G062200*). (**E**) Association between genotypes of the most significant SNP and ABD. (**F**) Changes in *PpLNK1* expression in three cultivars in a climate warming simulation experiment. dpt, days post treatment. (**G**) Comparison of *PpLNK1* expression between accessions sensitive and insensitive to global warming at blooming. ** represents *P* < 0.01. (**H**) Comparison of BD between wild type (WT) and *PpLNK1* over-expression (OE) *A. thaliana* lines. * indicates *P* < 0.05.

Next, we performed GWAS for ABD to identify genetic loci associated with responses to global warming (Supplementary Fig. S9B). This revealed a strong association peak on chromosome 8 (*P* < 7.28 × 10^−8^) (Fig. 6C) in an area harboring 14 candidate genes around peak association. This association was also located at overlap among GWAS signals of CR, BD and LFMM analysis. The most significant SNP was located in a region upstream of *Prupe*.*8G062200*, with genotype of TT showing sensitive to global warming and CC insensitive (Fig. 6D and 6E). *Prupe*.*8G062200* encodes a putative night light-inducible and clock-regulated 1 (LNK1) protein, and showed high expression levels at blooming. A homolog of this gene in *A. thaliana* is involved in regulation of the circadian clock, which regulates *COL1* genes at warm temperatures, and thus a potential regulator of flowering time (Mikkelsen and Thomashow 2009; Rugnone et al. 2013). A simulation experiment showed that *PpLNK1* was up-regulated by rising temperatures during heat accumulation, suggesting that *PpLNK1* may be up-regulated by temperature rise in spring (Fig. 6F). In addition, expression of *PpLNK1* in peach accessions that are sensitive to global warming was significantly higher than in those that are insensitive (Fig. 6G). Notably, over-expression of *PpLNK1* in model plant, *A. thaliana*, led to the early flowering (Fig. 6H). Moreover, several cis-elements associated with temperature and light responsiveness was identified (Supplementary Table S10). Therefore, we conclude that *PpLNK1* may play important roles in regulating annual circadian clock of flowering time as influenced by rising temperature in peach. *PpLNK1* is thus a plausible candidate gene for responses to global warming, but further work will be necessary to provide more direct evidence of its roles. Collectively, our comprehensive analyses detected genomic loci associated with responses to global warming, which can improve our understanding of the genetic architecture of plant adaptation to global climate change.

Long-term observation of BD enabled multi-year GWAS. We identified a total of 713 SNPs associated with BD (*P* < 7.28 × 10^−8^), including 483 temporary associations that were identified only in one year, 214 associations in at least two years, and 16 stable associations in more than five years, of which several overlapped with previous reported QTLs (Fan et al. 2010) (Supplementary Table S11). Among stable associations, a strong association peak within a small intergenic region (Pp06: 15,327,714∼15,354,080) on chromosome 6 was identified in eight years of GWAS, which can be further developed for marker-assisted selection.

## Conclusions

Plant genomes have been shaped by natural selection during the local adaptation to diverse environmental conditions. Peach provides an excellent model to investigate the genetic basis and mode of adaptation to climate change, thanks to its relatively small genome size (∼227.4Mb) and extensive climatic variation across its native range. We generated a large variation map for peach through sequencing of a climate-extensive panel of 263 peach landraces and wild relatives. Notably, we first detected the genetic basis of adaptation to high altitudes for fruit species, *P. mira* (TB group), and we found that genes involved in the biosynthesis of flavonoids *(PpCHS2*) and stomatal development (*PpEPF1*) may play important roles in overcoming strong UV-B radiation and hypoxia, respectively, on the Tibet Plateau. We discovered that high sugar content and yellow flesh of peach in drought regions were drought-induced adaptive evolution mediated by interactions between the abscisic acid pathway, *PpSPS1* and carotenoids. More than nine thousand genomic loci, associated with 51 specific climate variables, were identified. These included several hotspots associated with temperature and precipitation, as well as a SNP associated with cold hardiness. Integrative analyses of selective sweeps and GWEAS suggest that peach adaptation was generated and enhanced by standing selection on multi sites. Genomic loci underlying the local adaption of BD and CR were found to be two evolutionary adaptations to low and high latitude regions. In addition, through data collected over a 30-year period, we identified a candidate genetic locus associated with responses to global warming in plant species.

This study provides new insights into peach adaptation to its habits and how climate has shaped the genome of a perennial tree plant through natural selection. These results also provide a new resource for studies of peach evolutionary biology and breeding, especially with regard to enhancing stress-resistance.

## Methods

### Plant materials and sequencing

A total of 263 peach accessions were sampled from the NPGRC (National Peach Germplasm Repository of China), except the 45 *P. mira* accessions, which were sampled from the Tibet plateau. These accessions, collected from almost all the distribution regions of peach landraces and wild relatives, including seven major ecotypes. These accessions included 45 of *P. mira* Koehne, 4 of *P. davidiana* (Carr.) Franch., 2 of *P. kansuensis* Rehd., a single *P. potaninii* Batal., 205 of *P. persica* L., and 6 of *P. ferganensis* Kost. et Riab (Supplemental Table 1). Of these, *P. persica* L. and *P. ferganensis* Kost. et Riab accessions belong to landraces, while the others are wild relatives. Total genomic DNA was extracted from young leaves using the cetyltriethylammnonium bromide (CTAB) method (Murray and Thompson 1980). At least 4 μg of genomic DNA from each accession was used to construct pair-end sequencing libraries with insert sizes of approximately 300-bp or 500-bp following the manufacturer’s instructions (Illumina Inc.) (Supplemental Table 1). A total of >1 Gb of sequence data was generated for each accession from 49-bp, 90-bp, or 125-bp paired-end reads, using the Illumina GA or HiSeq 2500 platform (Illumina, San Diego, USA) (Supplemental Table S1).

### Read mapping and variation calling

Pair-end reads from each accession were mapped to the peach Lovell genome (release v2.0) using BWA (Li and Durbin 2009) (Version: 0.7.12) with the following parameters: bwa mem -t 4 - M -R. Read alignments were converted into the BAM format, sorted according mapping coordinates, and PCR duplicates removed using the Picard package (http://broadinstitute.github.io/picard/; Version: 1.136) with default parameters. The coverage and depth of sequence alignments were computed using the Genome Analysis Toolkit (GATK, version: 3.4-46; see URLs) DepthOfCoverage program (McKenna et al. 2010). The coverage and depth of each accession are detailed in Supplemental Table S1.

To accurately identify SNPs, the low-quality alignments (a mapping quality score <20) were filtered using SAMtools (Li et al. 2009). SNP detection was performed using GATK HaplotypeCaller, which identifies SNPs by local *de novo* assembly of haplotypes in an active region (Depristo et al. 2011). The detailed processes were as follows: (1) After filtering the low-quality alignments, the reads around the INDELs were realigned through two steps, including identifying regions where realignment was needed using the GATK RealignerTargetCreator package, and realigning the regions found in the first step GATK IndelRealigner package. Next, a realigned BAM file for each accession, which was used for SNP detection, was generated using GATK PrintReads packages. (2) SNPs were detected at a population level using the realigned BAM file with GATK HaplotypeCaller. To reduce the number of false positives, a high SNP confidence score was set with the following parameters: -stand_call_conf 30 -stand_emit_conf 40. (3) To ensure the quality of variant calling, a hard filter was applied for the raw SNPs with SNP quality > 40 and the number of supporting reads > 2, using GTAK VariantFiltration, with the following parameters: QUAL < 40, QD < 2.0, FS > 60.0, MQ < 40.0, MQRankSum < −12.5, ReadPosRankSum < −8.0, -cluster 3, -window 10.

The accuracy of SNPs was assessed using a Sequenom MassARRAY platform (Sequenom, San Diego, USA), following the manufacturer’s protocol. A total of 18 randomly selected SNPs was investigated in 130 accessions. The list of accessions is provided in Supplemental Table S2.

INDEL calling was performed using the same pipeline as the SNP calling since the GATK is capable of calling SNPs and INDELs simultaneously. To reduce the number of false positives, we also applied a harder filter for raw INDELs using GTAK VariantFiltration with the following parameters: QD < 2.0, FS > 200.0, ReadPosRankSum < −20.0. Insertions and deletions ≤6 bp were defined as the small INDELs.

SV calling was performed using the SpeedSeq (Chiang et al. 2015), DELLY (Tobias et al. 2012), and manta (Chen et al. 2016) programs. For SpeedSeq calling, paired-end reads were mapped to the reference genome using the ‘align’ module in SpeedSeq and the following parameters: speedseq align -R -t 4. Three BAM files were generated, including a full, duplicate-marked, sorted BAM, a BAM file containing split reads, and a BAM file containing discordant read-pairs. SVs were identified using the ‘sv’ module in SpeedSeq, using the following settings: speedseq sv -o -x -t 25 -R -B -D -S -g -P. For DELLY calling, mapped pair-end reads in BAM format, generated by BWA-MEM (Li and Durbin 2009) after sorting and marking PCR duplicates, were used as input files. SVs were identified using the call module in DELLY with default parameters. SV files in VCF format for all of 263 samples were merged into a population level VCF file using BCFtools (Li et al. 2009). For SV calling with manta, the same BAM files with DELLY were used to detect SVs, with default parameters. SV files for 263 accessions were then merged using SURIVAR (Jeffares et al 2017) and genotyped using SVtyper (Chiang et al. 2015) with default parameters. Finally, SVs identified by at least two callers were designated as the final set of SVs.

### SNP annotation

SNP annotation was performed based on genomic locations and predicted coding effects, according to the peach genome annotation (release annotation v2.1, see URLs), using the snpEff (Cingolani et al. 2012) (Version: 4.1g). The final SNPs were categorized in exonic regions, intronic regions, splicing sites, 5’ UTRs and 3’ UTRs, upstream and downstream regions, and intergenic regions, based on the peach genome annotation. SNPs in coding sequence were further grouped into synonymous SNPs (no amino acid changes) and nonsynonymous SNPs (amino acid changes). SNP effects were further divided into four types according to their impacts on gene function, including HIGH, MODERATE, LOW, and MODIFIER.

### Population genetics analysis

To build a phylogenetic tree, we selected a subset of 2,468,307 SNPs with minor allele frequency (MAF) >0.05 in all 263 accessions from the final SNP data set (4,611,842). A neighbor-joining tree was constructed using PHYLIP (Felsentein 1989) (Version:3.696) on the basis of the distance matrix with 1,000 bootstrap replicates. The software FigTree (http://tree.bio.ed.ac.uk/software/figtree/; version: 1.4.2) was used to visualize the neighbor-joining tree. The principal component analysis (PCA) was performed based on the same SNPs data set (2,468,307 SNPs with MAF > 0.05) using the smartpca program in the EIGENSFOT^70^ software (Version: 6.0.1) with default settings (Price et al. 2006). The first three eigenvectors were used to plot the data in two and three dimensions. The population structure was also investigated using the same SNP data set (2,468,307 SNPs with MAF>0.05) with the FRAPPE (Version: 1.1) software (Tang et al. 2005), which is based on a maximum likelihood method. We ran 10,000 iterations, and the numbers of clusters (*K*) were set from 2 to 8.

### Identification of select sweeps

To detect signals of selective sweeps, we selected three distinct genome-wide selection metrics for each group (excluding the TB group), including the reduction of nucleotide diversity (π), Tajima’s *D*, and genetic differentiation (*F*_ST_). We calculated these three selection metrics based on all SNPs (4,611,842) using VCFtools (Danecek et al. 2011) (Version: 0.1.13), with a 10-kb window and a step size of 1 kb. We defined the empirical top 5% of windows or regions as candidate selective outliers for each selection scan metric. The adjacent selective outliers were merged. For each population, selection outliers detected in at least two of the selection scan metrics were designated as the candidate selection regions (CSRs). The TB group consisted of wild relatives (*P. mira*) and three other methods were used to detect selective sweeps: Tajima’s *D*, RAiSD (Alachiotis et al. 2018), and CLR (Pavlidis et al. 2013). Similarly, the top 5% of windows or regions identified in at least two metrics were designated as candidate selective sweeps.

### Collection of climate variables

A total of 51 environmental variables were selected as being essential for peach growth and survival (Supplemental Table S6), representing extremes and seasonality of temperature and precipitation, altitude, latitude, relative air humidity, water vapor pressure, growing season lengths, and aridity. Of these, 39 datasets of climate variables were downloaded from WorldClim (http://www.worldclim.org; version: 1.4), with a resolution of 2.5 minutes, and climate variables for each accession were extracted using DIVA-GIS (http://www.diva-gis.org; version: 7.5) (Supplemental Table S6). Six climate variables were downloaded from CDMC (http://data.cma.cn/en/?r=site/index) and climate variables for each accession were extracted using ArcGIS (http://www.arcgis.com; version: 10.3) (Supplemental Table S6). Four climate variables were downloaded from the FAO (http://www.fao.org/geonetwork/srv/en/main.home), with a resolution of 5 minutes or 10 minutes and climate variables for each accession were extracted using ArcGIS (Supplemental Table S7). Altitude and latitude for each accession were recorded using a GPS (Magellangps triton 300E; http://www.magellangps.com) when the accessions were collected.

### Genome-wide environmental association study (GWEAS)

GWEAS was performed for 51 climate variables using 4,611,842 high-quality SNPs. The association analyses were performed using the mixed linear model (MLM) with Efficient Mixed-Model Association eXpedited (EMMAX) software (Zhou and Stephens 2012). To minimize the number of false positives and increase statistical power, population structure was corrected using a kinship matrix, which was estimated with EMMAX emmax-kin program (Zhou and Stephens 2012). The genome-wide significance thresholds of the GWEAS were determined using the Bonferroni test. Based on a nominal level of 0.05, the threshold was set as 0.05/total SNPs (log10(*P*) = −7.13).

### Functional enrichment and pathway analysis

To test whether candidate genes were overrepresented among lists from known biological processes, gene families and pathways, a functional enrichment and pathway analysis was performed based on Fisher exact tests (*P* < 0.05), using the Database for Annotation, Visualization and Integrated Discovery (DAVID) (Huang et al. 2009) (Version: 6.7). To obtain the comprehensive functional annotations, a list of annotation categories was selected, including GO terms and KEGG pathway. The annotation analysis was performed for genes that were in selective sweeps and GWEAS associations.

### Phenotyping and genome-wide association study (GWAS)

The first bloom date (BD) was measured at the National Peach Germplasm Repository of China (NPGRC) (N34.71°, E113.70°, A.S.L. 74 m), located in Zhengzhou, Henan Province, China. The first bloom date data used span February 25 to April 25 from 1983 to 2011 as this period captured the majority of diversity of BD. A total of 89 accessions, with each represented by two replicates, were used to investigate BD (Supplemental Fig. S9A). The first bloom date was defined as the day when approximately 5% of the flowers have completely opened. The advance in bloom date (ABD) for each accession was estimated using a linear regression analysis, based on the BD from 1983 to 2011. The ABD information for each accession is detailed in Supplemental Fig. S9B. To identify genetic loci associated with ABD, GWAS was performed for ABD based a set of 873,895 SNPs, identified after removing SNPs with low-frequency (MAF < 0.05) and a high missing rate (missing rate > 0.2) using the EMMAX program (Zhou and Stephens 2012). To minimize the number of false positives and to increase the statistical power, population structure was corrected using a kinship matrix, which was calculated with EMMAX emmax-kin program (Zhou and Stephens 2012). The genome-wide significance threshold of the GWAS was determined using the Bonferroni test. Based on a nominal level of 0.05, the threshold was set as 0.05/total SNPs (log10(*P*) = −7.08). GWAS was also performed for yearly BD data from 1983 to 2011 based on the same SNP data set, using the same method as above.

For CR, phenotyping analyses were performed in 2011 and 2012 as in Fan et al (2010). A 0-7.2°C model was chosen to evaluate CR and the number of hours in this range (chilling hours; CHs) was counted, starting when the daily average air temperature dropped to below 7.2°C. Starting at 50 CHs, the branches of each accession were cut every 50 CHs until 1,300 CHs. For each accession, two clones were sampled, and three branches longer than 40 cm with floral buds were taken from each clone. Branch cuttings were placed in water in a greenhouse at 25°C and a 16 h/8 h photoperiod to force floral bud break. The frequency of floral bud break was evaluated after two weeks. The CR of an accession was defined as being sufficient at a specific sampling time if 50% of floral buds on the branch cuttings opened. GWAS for CR was also performed using MLM in EMMAX.

Cold hardiness was evaluated using a conductance-based semi-lethal temperature method in December-January of 2013-2014 on 143 accessions. Six annual branches longer than 20 cm were sampled for each accession. A total of six cold treatments were used: −10, −15, −20, −25, −30, and −35°C. Branch cuttings were incubated in freezer with the six treatments for 24 h. After cold treatments, the cuttings were placed at 0°C for 8 h. Branch cuttings were then cut into 2 mm segments. A total of 2 g of segments was used to measure the conductance, with three biological replicates. The initial conductance (C1) was measured after a 12 h steep in 10 ml water. The final conductance (C2) was measured after boiling the samples for 20 min and leaving them to cool to room temperature for a subsequent 2 h period. The relative conductance (RC) was calculated using following formula:

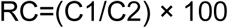

Finally, the semi-lethal temperature (LT50) was calculated using a logistic function based on RC.

### Yeast one-hybrid assay

Yeast one-hybridization assay was performed using the Matchmaker^®^ Gold Yeast One-Hybrid System (Clontech, Palo Alto, CA, USA). The promoter sequence (upstream 2kb from the start codon) of the sucrose phosphate synthase, *PpSPS1* (*Prupe*.*1G483200*), was cloned into the pAbAi vector. Similarly, the full-length of ABA-responsive element binding 1, *PpAREB1* (*Prupe*.*1G434500*), was subcloned into the pGADT7 AD vector. The auto-activation and TF– protein interaction analyses were conducted according to manufacturer’s protocol.

### Scanning electron microscopy (SEM)

Stomata were examined by SEM in young leaves from three accessions from the TB group and three accessions from the NP group, representing high-altitude and low-altitude accession, respectively. Three replicates were sampled from each accession. Samples were fixed in 2.5% glutaraldehyde (pH = 7.4) for 24 h at 4**°C**. Subsequently, fixed samples were dehydrated using an ethanol series (30% ethanol, 20 min; 50% ethanol, 20 min; 70% ethanol, 20 min; 100% ethanol, 30 min (twice)). The dehydrated samples were then dried in a critical-point drying apparatus (Quorum K850; England). Dried samples were mounted on stubs and sputter-coated with gold (FEI; America) and observed under a scanning electron microscopy (SEM) (FEI Quanta 250; America).

### RNA-Seq analysis

For drought stress treatment, four-year-old potted peach seedlings from peach cultivar “Dong Xue Mi Tao” were used. Fruit flesh were taken at six drought stress treatment time points, including 6 hours, 12 hours, 24 hours, 3 days, 6 days, and 12 days. For expression profiles in different tissues, roots, leaves, fruit, flowers, phloem, and seeds were sampled from “Aba Guang He Tao” (high-altitude) and “B-4” (low-altitude). For the expression of *PpCCD4*, fruit fleshes were sampled from “Zao Huang Pan Tao” (yellow-fleshed) and “Zhong Tao Hong Yu” (white-fleshed) at 10, 50, and 90 days post bloom date (dpb). For the expression of *PpSPS1*, fruit fleshes were sampled from “Chinese cling” at 20, 40, 60, 80, 100, 120 dpb. Three biological replicates were collected for each sample. The tissues were immediately frozen in liquid nitrogen and then ground to fine powder. Total RNA was extracted using a quick extraction kit (Aidlab, Beijing, China). First and second strand complementary DNA (cDNA) was synthesized using a cDNA Synthesis System kit (TOYOBO, Osaka, Japan), following the manufacturer’s protocol. Double-strand cDNAs were then purified and adapters were ligated to the short fragments. The constructed RNA-Seq libraries were sequenced using the Illumina HiSeq 2000 platform (Illumina, San Diego, USA) in paired-end 150-bp mode. Low-quality reads were filtered from the raw reads using Trimmomatic (Bolger et al. 2014). Data analysis followed the protocol proposed by Pertea et al (2016). Cleaned reads were mapped to the peach reference genome using Hisat2 (Version 2.0.5) (Kim et al. 2015) with default parameters. Transcript abundances were calculated and transcript assembly was performed using Stringtie (Pertea et al. 2015). DEG analysis was carried out using the R package ballgown (Frazee et al. 2015).

### Over-expression of candidate genes in *A. thaliana*

The full-length open reading frames of three peach genes, *PpEPF1* (*Prupe*.*3G235800*), *PpSVP* (*Prupe*.*8G069300*), and *PpLNK1* (*Prupe*.*8G062200*), were amplified by PCR using cDNAs derived from young leaves of “Aba Guang He Tao”, ‘Nanshan Tian Tao’ (CR=200h), and “Nanshan Tian Tao”, respectively. The PCR products were cloned into the pBI121 vector driven by the cauliflower mosaic virus (CaMV) 35S promoter at Sangon Biotech (Sangon, Shanghai, China). The resulting constructs were then transformed into *A. thaliana* Columbia type using *Agrobacterium tumefaciens* GV3101 and positive transformants selected with kanamycin. Ten transgenic lines for each gene were used to evaluate the flowering time. The stomata size and density were observed under a light microscope (Olympus BX51, Tokyo, Japan) with a 400 × objective lens.

### RNA extraction and expression analysis using qRT-PCR

For *PpSVP* expression analysis, floral buds from ‘Nanshan Tian Tao’ were sampled on October 15, November 15, December 15, January 15, February 15, March 15 in 2016-2017. *PpLNK1* expression was measured in floral buds (blooming soon) from three global warming-sensitive accessions (‘Wu Yue Xian’, ‘Nanshan Tian Tao’, and ‘Li He Pan Tao’) and three global warming-insensitive accessions (‘Xinjiang Pan Tao’, ‘Wuhan 2’, and ‘Kashi 2’) at 2016 and 2017. For *PpAHP5*, the phloem (including cambium) was collected from two cultivars ‘Hunchun Tao’ (cold resistant) and ‘Nanshan Tian Tao’ (cold sensitive) after 24 hours treatment under −28 °C refrigerator and following 21 °C incubation in water. For each sample, three biological replicates were used. Total RNA was extracted using an extraction kit (Aidlab, Beijing, China) and first-strand cDNA was synthesized with 1µg RNA using a FastQuant RT Kit (with gDNase) (TIANGEN, Beijing, China). Gene-specific primers were designed using Primer-BLAST software (National Center for Biotechnology Information, Maryland, USA). qRT-PCR was performed using a SYBR green I master kit (Roche Diagnostics, Indianapolis, USA) with the LightCycler System (Roche LightCycler 480, Indianapolis, USA), following the manufacturer’s protocol. Relative expression levels were calculated using the 2^-ΔΔCT^ method. A β-actin was used as the reference gene.

### Global warming simulation experiment

The global warming simulation experiment was performed in 2016-2017. Three peach cultivars (Nanshan Tian Tao, Hu Jing Mi Lu, and Maravila), each with two clones, were used as plant materials. For each cultivar, ∼30 annual branches longer than 40 cm with floral buds were taken from each clone when the winter chill accumulation was ∼900 chilling hours (0∼7.2°C, excluding 0°C). Branch cuttings were placed in water in greenhouse at 25°C and with a 16 h/8 h photoperiod, to simulate climate warming. The ratio of bud break was investigated daily, starting from the day that the branch cuttings were placed in the greenhouse. The floral buds, excluding the tegmentum, were collected weekly and frozen in liquid nitrogen. The sampled floral buds were used for qRT-PCR analyses following the protocol described above.

### Data access

Raw sequence data have been deposited in the NCBI Short Read Archive (SRA) under accession SRP108113. SNPs and SVs in Variant Call Format (VCF) have been deposited into the Figshare database (SNPs: https://figshare.com/articles/SNPs_for_263_peach_accessions/7636715, SVs: https://figshare.com/articles/SVs_for_peach_sequencing/7636721). All other relevant data are contained within the paper and available in supplementary files.

## Acknowledgements

This work was supported by grants from the Agricultural Science and Technology Innovation Program (CAAS-ASTIP-2020-ZFRI-01), the National Natural Science Foundation of China (31572094), the Crop Germplasm Resources Conservation Project (2016NWB041), and the US National Science Foundation (IOS-1339287 and IOS-1539831). We thank Prof. Jialong Yao from The Plant and Food Research Institute of New Zealand and Dr. Amandine Cornille from Université Paris-Sud for helpful suggestions in paper writing. We thank Dr. Yanling Wen from Beijing Institute of Genomics, Chinese Academy of Sciences for assistance in data visualization.

## Author contributions

L.W., S.H., Z.F. and W.G. designed and managed the project; Y.L., G.Z., X.Z., S.Z. and C.C. collected materials; Y.L., P.Z., J.G., X.W., and Q.Z. prepared and purified DNA samples; Y.L., K.C., and N.L. performed the data analyses; Y.L., T.D., J.W., L.G., Q.H., and W.F. performed phenotyping. Y.L. performed the molecular experiment. Y.L. and K. C. wrote the paper; L.W., Z.F., W.G., and S.H. revised the paper. All authors read and approved the final manuscript.

## References

Alachiotis N, Pavlidis P. 2018. RAiSD detects positive selection based on multiple signatures of a selective sweep and SNP vectors. Commun Biol 1: 79.

Blanquart F, Kaltz O, Nuismer SL, Gandon S. 2013. A practical guide to measuring local adaptation. Ecol Lett 16: 1195–1205.

Bolger A, Scossa F, Bolger ME, Lanz C, Maumus F, Tohge T, Quesneville H, Alseekh S, Sørensen I, Lichtenstein G, et al. 2014. The genome of the stress-tolerant wild tomato species Solanum pennellii. Nat Genet 46: 1034–1038.

Bolger AM, Lohse M, Usadel B. 2014. Trimmomatic: a flexible trimmer for Illumina sequence data. Bioinformatics 30: 2114–2120.

Cao K, Zheng Z, Wang L, Liu X, Zhu G, Fang W, Cheng S, Zeng P, Chen C, Wang X, et al. 2014. Comparative population genomics reveals the domestication history of the peach, *Prunus persica*, and human influences on perennial fruit crops. Genome Biol 15: 415.

Cao K, Zhou Z, Wang Q, Guo J, Zhao P, Zhu G, Fang W, Chen C, Wang X, Wang X, et al. 2016. Genome-wide association study of 12 agronomic traits in peach. Nat Commu 7: 13246.

Chen X, Schulz-Trieglaff O, Shaw R, Barnes B, Schlesinger F, Källberg M, Cox AJ, Kruglyak S, Saunders CT. 2016. Manta: rapid detection of structural variants and indels for germline and cancer sequencing applications. Bioinformatics 32: 1220–1222.

Chiang C, Layer RM, Faust GG, Lindberg MR, Rose DB, Garrison EP, Marth GT, Quinlan AR, Hall IM. 2015. SpeedSeq: ultra-fast personal genome analysis and interpretation. Nat Methods 12: 966–968.

Cingolani P, Platts A, Wang L, Coon M, Nguyen T, Wang L, Land SJ, Lu X, Ruden DM. 2012. A program for annotating and predicting the effects of single nucleotide polymorphisms, SnpEff. Fly 6: 80.

Danecek P, Auton A, Abecasis G, Albers CA, Banks E, DePristo MA, Handsaker RE, Lunter G, Marth GT, Sherry ST. 2011. The variant call format and VCFtools. Bioinformatics 27: 2156–2158.

Depristo MA, Banks E, Poplin R, Garimella KV, Maguire JR, Hartl C, Philippakis AA, del Angel G, Rivas MA, Hanna M, et al. 2011. A framework for variation discovery and genotyping using next-generation DNA sequencing data. Nat Genet 43: 491–498.

Falchi R, Vendramin E, Zanon L, Scalabrin S, Cipriani G, Verde I, Vizzotto G, Morgante M. 2013. Three distinct mutational mechanisms acting on a single gene underpin the origin of yellow flesh in peach. Plant J 76: 75–87.

Fan S, Bielenberg DG, Zhebentyayeva TN, Reighard GL, Okie WR, Holland D, Abbott AG. 2010. Mapping quantitative trait loci associated with chilling requirement, heat requirement and bloom date in peach (*Prunus persica*). New Phytol 185: 917–930.

Felsenstein J. 1989. PHYLIP-phylogeny inference package (version 3.2). Cladistics 5: 164–166.

Ferrer JL, Jez JM, Bowman ME, Dixon RA, Noel JP. 1999. Structure of chalcone synthase and the molecular basis of plant polyketide pathway. Nat Struct Biol 6: 775–784.

Fournier-Level A, Korte A, Cooper MD, Nordborg M, Schmitt J, Wilczek AM. 2011. A map of local adaptation in *Arabidopsis thaliana*. Science 334: 83–86.

Frazee AC, Pertea G, Jaffe AE, Langmead B, Salzberg SL, Leek JT. 2015. Ballgown bridges the gap between transcriptome assembly and expression analysis. Nat Biotech 33: 243–246.

Hancock AM, Brachi B, Faure N, Horton MW, Jarymowycz LB, Sperone FG, Toomajian C, Roux F, Bergelson J. 2011. Adaptation to climate across the *Arabidopsis thaliana* genome. Science 334: 83–86.

Hara K, Yokoo T, Kajita R, Onishi T, Yahata S, Peterson KM, Torii KU, Kakimoto T. 2009. Epidermal cell density is autoregulated via a secretory peptide, EPIDERMAL PATTERNING FACTOR 2 in *Arabidopsis* leaves. Plant Cell Physiol 50: 1019–1031.

Harter RD. 1983. Effect of soil pH on adsorption of lead, copper, zinc, and nickel. Soil Sci Soc Am J 47: 47–51.

Hu Y, Jiang L, Wang W, Yu D. 2013. Jasmonate regulates the INDUCER OF CBF EXPRESSION–C-REPEAT BINDING FACTOR/DRE BINDING FACTOR1 Cascade and freezing tolerance in *Arabidopsis*. Plant Cell 25: 2907–2924.

Huang DW, Sherman BT, Lempicki RA. 2009. Systematic and integrative analysis of large gene lists using DAVID Bioinformatics Resources. Nat Protoc 4: 44–57.

Jeffares DC, Jolly C, Hoti M, Speed D, Shaw L, Rallis C, Balloux F, Dessimoz C, Bähler J, Sedlazeck FJ. 2017. Transient structural variations have strong effects on quantitative traits and reproductive isolation in fission yeast. Nat Commun 8: 14061.

Jeon J, Kim J. 2013. *Arabidopsis* response regulator1 and Arabidopsis histidine phosphotransfer protein2 (AHP2), AHP3, and AHP5 function in cold signaling. Plant Physiol 161: 408–424.

Kim D, Langmead B, Salzberg SL. 2015. HISAT: a fast spliced aligner with low memory requirements. Nat Methods 12: 357–360.

Kobashi K, Gemma H, Iwahori S. 2000. Abscisic acid content and sugar metabolism of peaches grown under water stress. J Amer Soc Hort Sci 125: 425–428.

Kobashi K, Sugaya S, Gemma H, Iwahori S. 2001. Effect of abscisic acid (ABA) on sugar accumulation in the flesh tissue of peach fruit at the start of the maturation stage. Plant Growth Regul 35: 215–223.

Lasky JR, Upadhyaya HD, Ramu P, Deshpande S, Hash CT, Bonnette J, Juenger TE, Hyma K, Acharya C, Mitchell SE. 2015. Genome-environment associations in sorghum landraces predict adaptive traits. Sci Adv 1: e1400218.

Lastdrager J, Hanson J, Smeekens S. 2014. Sugar signals and the control of plant growth and development. J Exp Bot 65: 799–807.

Li H, Durbin R. 2009. Fast and accurate short read alignment with Burrows-Wheeler transform. Bioinformatics 25: 1754–1760.

Li H, Durbin R. 2011. Inference of human population history from individual whole-genome sequences. Nature 475: 493–496.

Li H, Handsaker B, Wysoker A, Fennell T, Ruan J, Homer N, Marth G, Abecasis G, Durbin R, 1000 Genome Project Data Processing Subgroup. 2009. The sequence alignment/map format and SAMtools. Bioinformatics 25: 2078–2079.

Li J, Oulee TM, Raba R, Amundson RG, Last RL. 1993. *Arabidopsis* Flavonoid Mutants Are Hypersensitive to UV-B Irradiation. Plant Cell 5: 71–179.

Li Y, Cao K, Zhu G, Fang W, Chen C, Wang X, Zhao P, Guo J, Ding T, Guan L, et al. 2019. Genomic analyses of an extensive collection of wild and cultivated accessions provide new insights into peach breeding history. Genome Biol 20(1): 36.

Li Y, Wang L, Zhu G, Fang W, Cao K, Chen C, Wang X, Wang, X. (2016). Phenological response of peach to climate change exhibits a relatively dramatic trend in China, 1983-2012. Sci Hortic-Amsterda 209:192–200.

Li Z, Reighard GL, Abbott AG, Bielenberg DG. 2009. Dormancy-associated MADS genes from the *EVG* locus of peach [*Prunus persica* (L.) Batsch] have distinct seasonal and photoperiodic expression patterns. J Exp Bot 60: 3521–3530.

Ma Q, Sun M, Lu J, Liu Y, Hu D, Hao Y. 2017. Transcription factor AREB2 is involved in soluble sugar accumulation by activating sugar transporter and amylase genes. Plant Physio 174: 2348–2362.

Martínez-García PJ, Parfitt DE, Ogundiwin EA, Fass J, Chan HM, Ahmad R, Lurie S, Dandekar A, Gradziel TM, Crisosto CH. 2013. High density SNP mapping and QTL analysis for fruit quality characteristics in peach (*Prunus persica* L.). Tree Genet & Genomes 9: 9–36.

McKenna A, Hanna M, Banks E, Sivachenko A, Cibulskis K, Kernytsky A, Garimella K, Altshuler D, Gabriel S, Daly M, et al. 2010. The Genome Analysis Toolkit: a MapReduce framework for analyzing next-generation DNA sequencing data. Genome Research 20: 1297–1303.

Menzel A, Sparks TH, Estrella N, Koch E, Aasa A, Ahas P, Alm-Kubler K, Bissolli P, Braslavska O, Briede A, et al. 2006. European phenological response to climate change matches the warming pattern. Glob Chang Biol 12: 1969–1976.

Mikkelsen MD, Thomashow MF. 2009. A role for circadian evening elements in cold-regulated gene expression in *Arabidopsis*. Plant J 60: 328–339.

Monihan SM, Magness CA, Yadegari R, Smith SE, Schumaker KS. 2016. Arabidopsis CALCINEURIN B-LIKE10 functions independently of the SOS pathway during reproductive development in saline conditions. Plant Physio 171: 369–379.

Murray M, Thompson WF. 1980. Rapid isolation of high molecular weight plant DNA. Nucleic Acids Res 8: 4321–4326.

Pertea M, Kim D, Pertea G, Leek JT, Salzberg SL. 2016. Transcript-level expression analysis of RNA-seq experiments with HISAT, StringTie and Ballgown. Nat Protoc 11: 1650–1667.

Pertea ML, Pertea GM, Antonescu CM, Chang TC, Mendell JT, Salzberg SL. 2015. StringTie enables improved reconstruction of a transcriptome from RNA-seq reads. Nat Biotech 33: 290–295.

Price AL, Patterson NJ, Plenge RM, Weinblatt ME, Shadick NA, Reich D. 2006. Principal components analysis corrects for stratification in genome-wide association studies. Na Genet 38: 904–909.

Pritchard J, Di Rienzo A. 2010. Adaptation-not by sweeps only. Nat Rev Genet 11: 665–667.

Rugnone ML, Faigón Soverna A, Sanchez SE, Schlaen RG, Hernando CE, Seymour DK, Mancini E, Chernomoretz A, Weigel D, Más P, et al. 2013. *LNK* genes integrate light and clock signaling networks at the core of the *Arabidopsis* oscillator. Proc Natl Acad Sci U S A 110: 12120–12125.

Sasaki R, Yamane H, Ooka T, Jotatsu H, Kitamura Y, Akagi T, Tao R. 2011. Functional and expressional analyses of *PmDAM* genes associated with endodormancy in Japanese apricot. Plant Physiol 157: 485–497.

Seguel A, Cumming JR, Klugh-Stewart K, Cornejo P, Borie F. 2013. The role of arbuscular mycorrhizas in decreasing aluminium phytotoxicity in acidic soils: a review. Mycorrhiza 23: 167–183.

Tang H, Peng J, Wang P, Risch N. 2005. Estimation of individual admixture: analytical and study design considerations. Genet Epidemiol 28: 289–301.

Tim W, Braun JV. 2013. Climate change impacts on global food security. Science 341: 508–513.

Tobias R, Zichner T, Schlattl A, Stütz AM, Benes V, Korbel JO. 2012. Delly: structural variant discovery by integrated paired-end and split-read analysis. Bioinformatics 28: i333–i339.

Verde I, Abbott AG, Scalabrin S, Jung S, Shu S, Marroni F, Zhebentyayeva T, Dettori MT, Grimwood J, Cattonaro F, et al. 2013. The high-quality draft genome of peach (*Prunus persica*) identifies unique patterns of genetic diversity, domestication and genome evolution. Nat Genet 45: 487–494.

Wang J, Ding J, Tan B, Robinson KM, Michelson IH, Johansson A, Nystedt B, Scofield DG, Nilsson O, Jansson S, Street NR, et al. 2018. A major locus controls local adaptation and adaptive life history variation in a perennial plant. Genome Bio 19: 72.

Wang L, Zhu GR, Fang WC. 2012. Peach genetic resource in China. China Agriculture Press.

Yan W, Liu H, Zhou X, Li Q, Zhang J, Lu L, Liu T, Liu H, Zhang C, Zhang Z, et al. 2013. Natural variation in *Ghd7.1* plays an important role in grain yield and adaptation in rice. Cell Res 23(7): 969–971.

Zhang Q, Chen W, Sun L, Zhao F, Huang B, Yang W, Tao Y, Wang J, Yuan Z, Fan G, et al. (2012). The genome of *Prunus mume*. Nat Commun 3: 1318.

Zheng Y, Crawford GW, Chen X. 2014. Archaeological evidence for peach (*Prunus persica*) cultivation and domestication in China. PLoS ONE 9: e106595.

Zhou X, Stephens M. 2012. Genome-wide efficient mixed-model analysis for association studies. Nat Genet 44: 821–824.

